# A novel biparatopic antibody-ACE2 fusion that blocks SARS-CoV-2 infection: implications for therapy

**DOI:** 10.1101/2020.06.14.147868

**Authors:** Xiaoniu Miao, Yi Luo, Xi Huang, Suki M. Y. Lee, Zhijun Yuan, Yongzhou Tang, Liandi Chen, Chao Wang, Wenchao Jiang, Wei Gao, Xuedong Song, Yao Yan, Tuling Pang, Yuefeng Zou, Weihui Fu, Liping Wan, Javier Gilbert-Jaramillo, Michael Knight, Tiong Kit Tan, Pramila Rijal, Alain Townsend, Joanne Sun, Xiaolin Liu, William James, Andy Tsun, Yingda Xu

## Abstract

In the absence of a proven effective vaccine preventing infection by SARS-CoV-2, or a proven drug to treat COVID-19, the positive results of passive immune therapy using convalescent serum provides a strong lead. We have developed a new class of tetravalent, biparatopic therapy, 89C8-ACE2. It combines the specificity of a monoclonal antibody (89C8) that recognizes the relatively conserved N-terminal domain (NTD) of the viral S glycoprotein, and the ectodomain of ACE2, which binds to the receptor-binding domain (RBD) of S. This molecule shows exceptional performance in vitro, inhibiting the interaction of recombinant S1 to ACE2 and transduction of ACE2-overexpressing cells by S-pseudotyped lentivirus with IC50s substantially below 100 pM, and with potency approximately 100-fold greater than ACE2-Fc itself. Moreover, 89C8-ACE2 was able to neutralize authentic virus infection in a standard assay at low nanomolar concentrations, making this class of molecule a promising lead for therapeutic applications.

## Introduction

The COVID-19 pandemic is the biggest global health threat in this decade. As of May 2020, studies on at least 120 possible COVID-19 vaccines were already underway, but whether a safe and effective product could be developed was still unclear. Looking back at history, scientists have never been able to develop a medically proven vaccine against any strain of coronavirus. The natural immunity to coronaviruses seem short-lived, previous studies suggested the possibility of re-infection after initial exposure to SARS-CoV^1^, ^2^. However, even if an effective vaccine proved possible, it may still require years to develop into a product. Thus, especially during a novel outbreak, there is an urgent need for the discovery of other therapeutic options. The administration of convalescent plasma collected from recovered patients has shown adequate safety and provided protection from clinical complications including fever and respiratory symptoms in severe COVID-19 patients^3^, ^4^. However, the success of convalescent plasma therapy hinges on many factors, including the availability of donors, concentration of anti-SARS-CoV-2 neutralizing antibodies, and the safe preparation of serum to eliminate risks of viral transmission via transfusion. During disease outbreaks, approaches to isolate the neutralizing antibodies from recovered patients have proved successful. Similar campaigns from industry (Eli Lilly & Co, Regeneron) and academia (Tsinghua University, Peking University, Oxford University) to discover and develop neutralizing antibodies (nAbs) have been launched for COVID-19 since the beginning of 2020. The monovalent affinity of these neutralizing antibodies towards the viral spike protein (S1) are usually relatively weak. This has made it necessary to screen for large numbers of antibodies to select for potent neutralizing antibodies, or apply computational approaches for *in silico* affinity maturation to improve binding affinities. To boost neutralization activity, a cocktail approach of combining two or more neutralizing antibodies has been sought but at the cost of increased manufacturing burden. An alternative approach to block viral infection is to target the viral entry receptor, ACE2. Recombinant ACE2 has been shown to reduce viral infection and growth in cell cultures including organoids by acting as a decoy for SARS-CoV-2^5^. APN01, a homodimer of ACE2, fails to effectively block the viral infection likely due to its dissimilar mode of interaction with viral S1 compared with its natural form and short serum half-life. The fusion of ACE2 to an Fc domain, producing a recombinant bivalent ACE2, could extend its physiological half-life and offer higher avid affinity towards viral S1, and thus increase the potency of blocking viral entry^6^.Here we describe the design and discovery of a biparatopic construct, in which a neutralizing antibody (89C8) that binds to the N-terminal domain (NTD) of S1 is fused to recombinant ACE2 (89C8-ACE2). 89C8-ACE2 offers superior binding affinity to viral S1 protein with potent neutralizing activity as demonstrated by pseudotype and native virus infectivity assays. This design may also offer neutralizing capacity towards different strains of coronaviruses by avoiding the potential loss of binding due to mutations in the receptor binding domain (RBD) of S1 protein, and offers insight into a universal therapeutic design that could be adopted for the treatment of other infectious diseases.

### Antibody selection

In this study, we aimed to isolate Abs against SARS-CoV-2. We first collected human peripheral venous blood samples from ten donors at the Fifth Affiliated Hospital, Sun Yat-Sen University. Recombinant S1 protein was used as bait to enrich S protein-specific memory B cells from the peripheral blood mononuclear cells (PBMCs) of COVID-19-recovered patients. Serum samples from 3 out of 10 donors displayed a strong reaction to SARS-CoV-2 S protein compared with equivalent samples obtained from healthy donor controls, as determined by biolayer interferometry (BLI) (Supplemental Figure 1), and corresponding B cells were subsequently used for the construction of libraries for yeast display screening. Three libraries (>10^8^ unique sequences each) of individual donors were constructed separately to minimize heavy/light-chain mispairing. S1-specific Fabs that were displayed on the yeast cells were selected using S1-protein-coated magnetic beads and subsequently sorted by FACS. A schematic diagram showing this workflow is illustrated in Figure 1.

**Figure 1.**
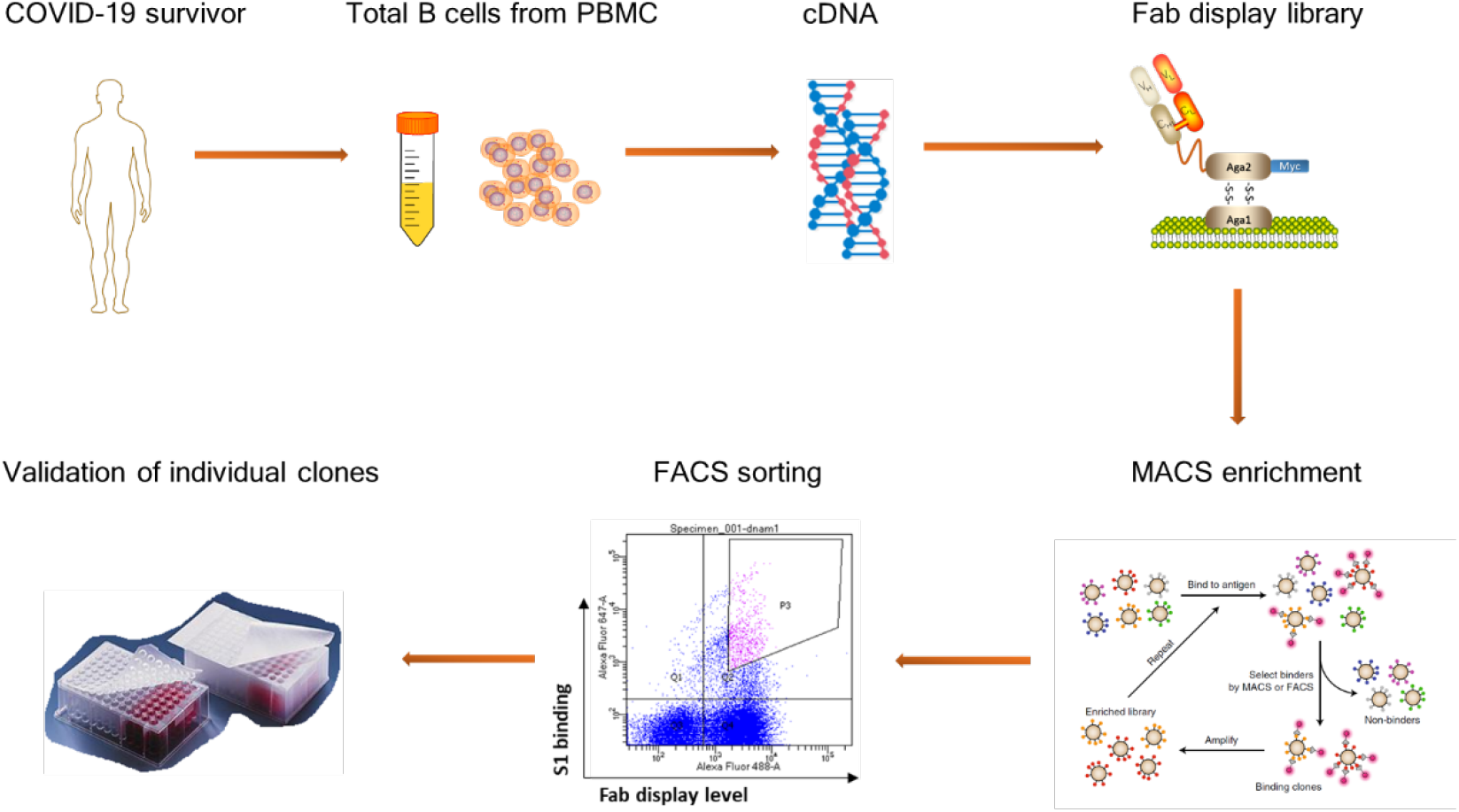
A schematic diagram showing the workflow of neutralizing antibody discovery

473 individual clones were picked for sequencing and a total of 115 unique, paired Fab sequences were obtained. 50 unique Fab sequences were sub-cloned into a eukaryotic expression vector for the generation of monoclonal antibody (mAb) protein for subsequent testing. These 50 antibodies were tested for binding to the HEK293 cells expressing the full length S1 protein of SARS-CoV-2 with follow-up characterization. Further considerations of lead selection included thermal stability, non-specific off-target binding and a faster intrinsic association constant towards S1 protein.

### Design and construction of the Biparatopic molecule

Next, our goal was to produce an anti-S1-recombinant ACE-2 fusion protein with biparatopic properties to provide superior binding affinity towards S1. Thus, we tested whether our antibodies could block the interaction between S1 and ACE2, with preference for the screening of non-blocking antibodies. One candidate, named 89C8, was chosen as the lead due to its faster association constant, clean binding towards untransfected HEK293 cells, and a superior Fab Tm (82 °C by DSF).

A biparatopic molecule was engineered with ACE2 fused with a stable (G4S)G linker to the heavy-chain C-terminal domain of 89C8 (Figure 2a). Alternative constructs with ACE2 fused to the N-terminus of either the LC or HC were also generated and included for comparison. We examined the binding of C-terminal and N-terminal ACE2 constructs to SARS-CoV-2 S1 in an Octet-based binding assay. Interestingly, only the C-terminal constructs showed strong binding, whereas none of the N-terminal constructs could show any binding to viral S1. 89C8 alone showed fairly strong monovalent binding to S1 with a relatively slow dissociation rate of ∼2E-04 S^-1^ (Figure 2b). ACE2-Fc exhibited a fast on/fast off profile with a monovalent binding affinity of ∼50 nM (Figure 2c). For the biparatopic molecule 89C8-ACE2, the monovalent dissociation was slower by a factor of 10 (Figure 2d). To determine the avidity of 89C8-ACE2, recombinant S1 protein was first biotinylated and immobilized on a streptavidin biosensor, then tested binding against 89C8-ACE2. The biparatopic molecule tightly bound to S1 with a strong multivalent affinity (Figure 2e). Compared to the monovalent Ab molecule, our multivalent design may provide stronger binding affinity and possess potential for longer-lasting protection from infection.

**Figure 2.**
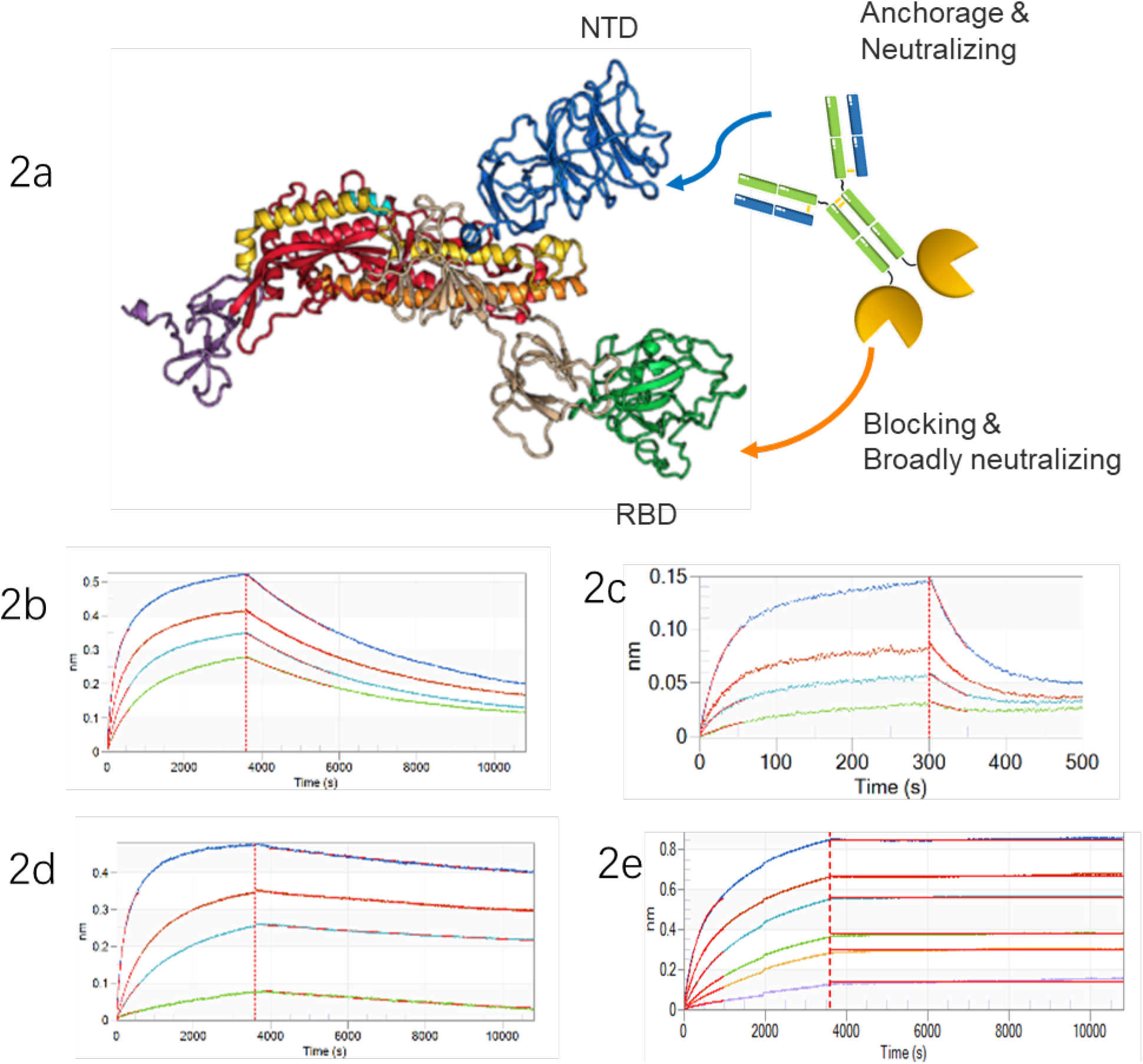
2a) schematic illustration of the biparatopic 89C8-ACE2 antibody-fusion design; 2b,c,d) BLI binding kinetics of 89C8, ACE2-Fc and 89C8-ACE2 binding to the spike protein S1 (monovalent K_D_); 2e) 89C8-ACE2 binding to the spike protein in the avid format.

To identify the site of the SARS-CoV-2 S protein bound by 89C8, a binding assay was performed using recombinant NTD and RBD of SARS-CoV-2 S1. 89C8 binds specially to the NTD but not RBD of S1 protein (Supplementary Figure 2), explaining the absence of competition between 89C8 and ACE2 for binding to SARS-CoV-2 S protein. ACE2-Fc fusion protein is a natural blocker of S1/ACE2 interaction, but its blocking activity may be relatively weak due to its low monovalent binding affinity. Cell-based blocking assay was used to investigate the blocking activity of 89C8-ACE2 (Figure 3a). 89C8 mAb and ACE2-Fc were also included as controls for comparison. ACE2-Fc was able to block the binding of soluble SARS-CoV-2 S1 with ACE2 receptor transiently expressed on CHO cells with an IC50 of 5nM. Compared to ACE2-Fc, 89C8-ACE2 enhanced the potency over a 100-fold with an IC50 of 55pM, whereas 89C8 did not have such blocking activity.

**Figure 3.**
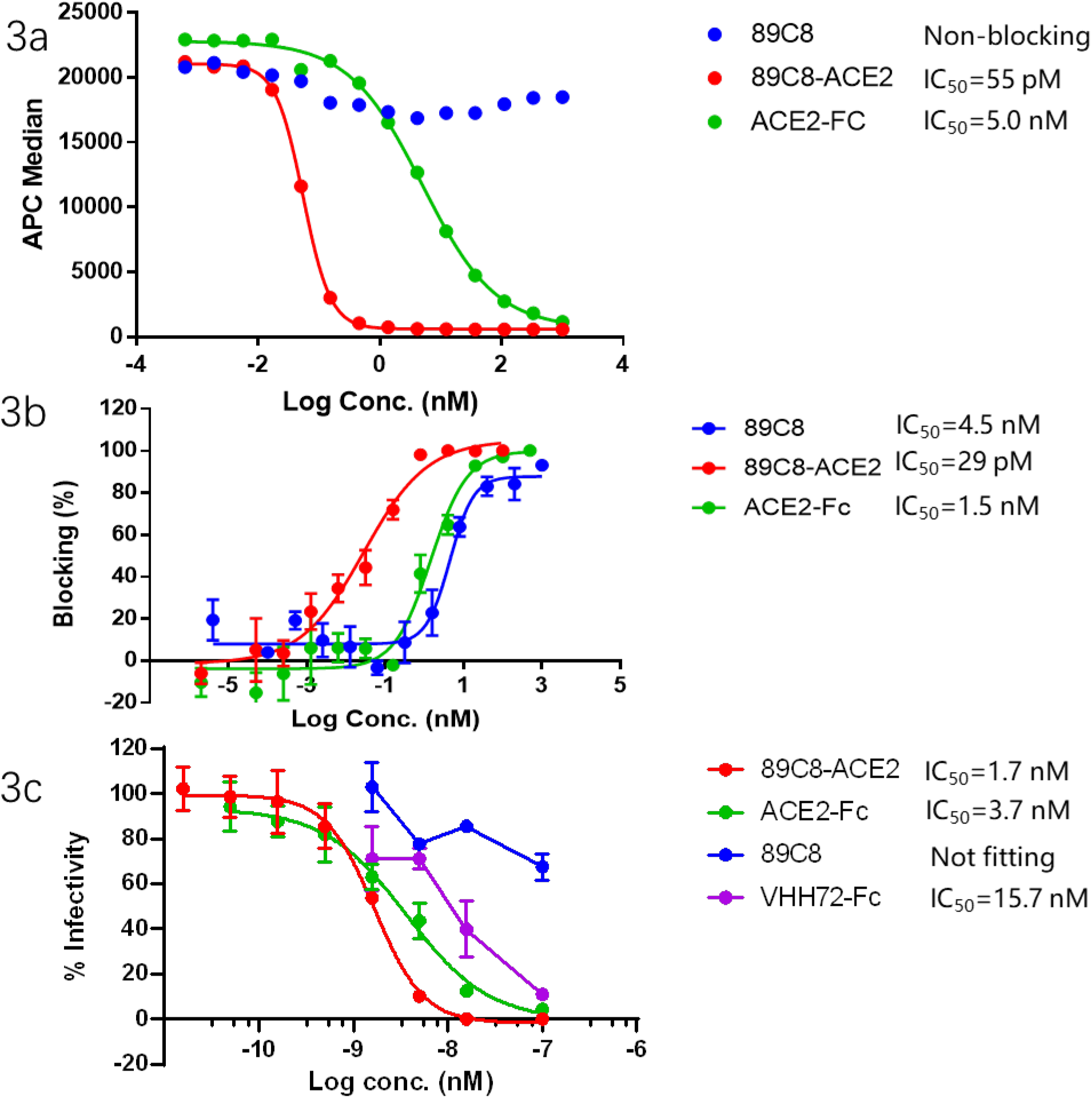
Blocking and neutralization experiments by 89C8, ACE2-Fc and 89C8-ACE2 between 3a) Spike protein S1 (m-Fc format) with CHO cells overexpressing ACE2; 3b) pseudovirus with HEK293 cells overexpressing ACE2; 3c) Live virus (Australia/Vic1/2020) plaque reduction neutralization test with Vero cells.

### Biparatopic 898-ACE2 is a potent neutralizer against SARS-CoV-2 infection

The neutralization potency of 89C8-ACE2 against SARS-CoV-2 infection was first assessed using an S-pseudotyped lentiviral reporter neutralization assay (Figure 3b). hACE2 overexpressing HEK293T cells were challenged with pseudotype virus encoding a luciferase reporter gene. Both 89C8-ACE2 and ACE2-Fc neutralized 100% of the input pseudotype virus, whereas 89C8 alone reached no more than 90% neutralization. Reciprocal IC50 neutralization titers (i.e. the concentration resulting in 50% reduction of infectivity) for 89C8-ACE2 against the pseudovirus was found to be 29 pM, which is much more potent compared to ACE2-Fc and 89C8, with IC50 values of 1.5 nM and 4.5 nM, respectively.

In the presence of 89C8-ACE2, infection of VeroE6 cells with authentic SARS-CoV-2 (Figure 3c) was neutralized with a PRNT50 and PRNT90 of 1.69 nM (95% confidence limits, 1.35-2.1 nM) and 6.23 nM (95 confidence limits 3.85-10.9 nM), respectively in a plaque reduction neutralization test. 89C8-ACE2 was superior compared to its parental molecule ACE2-Fc, which had PRNT 50 and PRNT 90 of 3.66 nM (95% confidence limits 2.28-6.05 nM) and 33.3 nM (95% confidence limits13.9-96 nM), respectively. However, 89C8 only showed very weak neutralization activity, below the confidence level for deducing a reliable PRNT50 value. Intriguingly, the neutralizing potential of 89C8-ACE2 was substantially higher than that of a potent neutralizing nanobody fusion, VHH72-Fc (15.7 nM), that binds to RBD-S1 with broad neutralization activity against betacoronaviruses^7^, revealing the superior neutralizing activity of 89C8-ACE2. ACE2-Fc showed stronger than expected neutralization activity, likely due to the presence of aggregates (∼40%).

## Discussion

COVID-19 is a novel and rapidly evolving pandemic causing severe morbidity & mortality and huge economic losses worldwide. The winter season has been correlated with the outbreak of SARS-CoV and SARS-CoV-2^8^ and it is possible that COVID-19 may reemerge in the Northern hemisphere during the winter. Presently, there have been no approved vaccines or targeted therapeutics available for COVID-19, and researchers are exploring various medical interventions including neutralizing antibodies. Development of nAbs targeting the vulnerable sites on viral surface proteins is considered to be an effective approach to combat viral infection, including coronaviruses^9^.

Potent nAbs to SARS and MERS often target S1 and S2 to disable binding between the RBD and receptor (targeting S1-RBD), interfere with the conformational change of S protein (targeting S1-NTD) or hinder S2-mediated membrane fusion (targeting S2)^10^, thus blocking CoV infection into host cells. Although S1-RBD is the most targeted domain for nAb development for SARS and COVID-19 (as RBD-specific antibodies have shown greater potency to neutralize infection with divergent virus strains), S1-RBD mutations with enhancing structural stability and ACE2 affinity were observed recently from circulating SARS-CoV-2 strains^11^. On the other hand, S1-NTD specific neutralizing antibodies have shown to be potent in blocking SARS-CoV infection^12^, due to the high homology of NTD in SARS-CoV and SARS-CoV-2, and thus the NTD could be a common target for vulnerability^13^.ACE2 is the cell entry receptor for SARS-CoV-2. Recently, studies have demonstrated that soluble ACE2 fused to Ig^14^ or human recombinant soluble ACE2^5^ can inhibit SARS-CoV-2 infection, suggesting that an ACE2 decoy could be a promising strategy for COVID-19 treatment. Previous studies have also demonstrated that ACE2 is also playing a protective role during pulmonary edema, so ACE2 serves both as the entry receptor for SARS-CoV and to protect the lung from injury^5^, ^15^. The severity of SARS could be partially explained by viral S binding to ACE2 at a site that does not interfere with its catalytic activity^16^, which then leads to endocytosis of the virus and loss of ACE2^17^, causing a detrimental cycle of viral infection and loss of protection from local lung injury. Therefore, an ACE decoy which lures the virus to attach itself to the decoy rather than the target host cells can protect the host from lung injury.

Here, we designed and generated a novel biparatopic antibody combining the above mentioned concepts by fusing two copies of ACE2 to the C-terminal domain of a neutralizing antibody (89C8) that targets S1-NTD. This design takes advantage of distinct neutralization mechanisms, via direct blocking and conformation-locking on the same S1 protein, and offers ultrahigh avidity. It was noted that 89C8-ACE2 binds to S1 with potent multivalent affinity and exhibits long-lasting effects. No detectable dissociation can be observed for up to 2 hours. Furthermore, we confirmed the high blocking potency of 89C8-ACE2 towards SARS-CoV-2 S1 and pseudovirus-expressing S Ag to hACE2-expressing cells, with IC50s in the picomolar range. 89C8-ACE2 exhibits substantial neutralizing activities against pseudovirus expressing S Ag or live virus *in vitro*, with superior activity over its parental mAb or ACE2 decoy as well as a known potent neutralizer, VHH72-Fc. VHH72-Fc possess a broad range of neutralizing activity against beta-coronaviruses. Our data here suggests that 89C8-ACE2 inhibits SARS-CoV-2 infection via a novel mechanism which takes advantage of added avidity effects via binding to both NTD-S1 and RBD-S1 simultaneously.

In our authentic virus neutralization test we designed the experiment to mimic viral infection and antibody/drug neutralization similar to a physiological setting, where long-lasting protection is required. In other live virus blocking/neutralizing assays, the virus and drug complexes are allowed to interact with the cells for just one hour before washing, followed by a short 8-hour incubation. This may not reflect the real neutralizing power of the drug. The experimental differences also reveals the weak neutralizing activity of 89C8 alone, or in general NTD-targeting antibodies. These antibodies neutralize viral infection through locking of the NTD conformation, thus preventing downstream events from occurring. As these antibodies do not prevent the association between the S1 RBD and cell surface ACE2, the antibody:NTD complex needs to be maintained tightly. On the other hand, for direct blockers, even after antibody dissociation, the S1 protein still requires virus particles to migrate into close proximity to ACE2-expressing cells, allowing formation of new S1:drug complex with circulating free drugs in serum. This maintains the neutralization complexes distal from ACE-2-expressing tissues.

In summary, our biparatopic design, either alone or in combination, may offer the potential to treat COVID-19, or can be adopted for other emerging diseases. Antibody combinations targeting distinct epitopes may act synergistically resulting in reduction of the dosage required for therapeutic potency as well as the risk of immune escape through mutational changes.

## Online Methods

### Yeast display library construction and antibody selection

Human peripheral blood samples were collected from recovered COVID-19 patients at the Fifth Affiliated Hospital, Sun Yat-Sen University. Mononuclear cells were isolated from the blood by Ficoll-Paque (GE Healthcare) density gradient centrifugation according to the manufacturer’s instructions. Total cellular RNA was isolated from cells using TRIZOL reagent (Invitrogen). cDNA was synthesized using SuperScript™ IV First-Strand Synthesis System (Invitrogen) primed with oligo (dT). Sequences of the antibody variable region were amplified using PCR. The light chain library was constructed by co-transforming linearized pFabVk vector and light chain PCR product by yeast gap repair. The Fab library was then constructed by co-transforming linearized pFabVH vector and heavy chain PCR product into the light chain library. Dual selection using Ura- and Trp-was performed to enrich the positive clones. Expression of Fab on the cell surface was induced in G/RCAA (substituting 2 g/L each of galactose and raffinose for the dextrose in SDCAA medium), and Fab-expressing cells were sorted by MACS (magnetic bead aided cell sorting, Mltenyi, Germany) then 4X by FACS (BD, AriaIII) using biotinylated SARS-CoV2 spike 1 protein.

### Recombinant antibody expression

The cDNAs encoding the variable regions of heavy (human IgG1 heavy chain) and light (kappa light chain constant regions) chains were cloned into expression plasmids (pCDNA3.1). The heavy- and light-chain encoding plasmids were transiently co-transfected into Expi293 cells (Life technologies) using PEI. Transfected cells were incubated for 7 days at 37 °C. IgG in the supernatants were purified for by Protein A magnetic beads (Genscript) according to manufacturer’s instructions.

### K_D_ determination

Binding affinities of IgGs (including 89C-ACE2 and ACE2-Fc) and S1 protein were determined by biolayer interferometry (BLI) using an Octet 384 (Pall Life Sciences). IgGs were loaded onto AHC sensors prior to exposure to SARS-CoV-2 S1 antigen in solution (300s or 3600s) for association and then dissociation in a buffer solution (300s or 7200s). Data acquisition was performed at 30 °C. Data was analyzed using the ForteBio Data Analysis Software. The data was fit to a 1:1 binding model to calculate an association and dissociation rate, and KD was calculated using the ratio kd/ka.

### Cell line generation and culture

HEK293T cells purchased from ATCC (CRL-3216TM) were cultured in DMEM with 10% FBS supplemented with GlutaMAX. Human ACE2 coding sequence (Uniport: Q9BYF1) was amplified (Genewiz) and cloned into pLVX-Puro vector. HEK293T cells were transfected with pLVX-Puro-hACE2 in addition to psPAX2 and pMD using Lipofectamine 3000 (Invitrogen). 24h post transfection, media was replaced with complete media and further incubated 24h before collection of supernatant. HKE293T or CHO-S were then infected with viral supernatant and selected using puromycin.

### Cell based blocking test

Gradient diluted antibodies or antibody-ACE2 fusion proteins were first pre-incubated with 0.1nM S1-mFc (SinoBiological) at 37 °C overnight, followed by incubation with CHO cells stably expressing ACE2 protein (CHO-ACE2) for 1h at RT. The cells were washed and subsequently incubated with APC-labeled goat anti-mouse IgG (ebioscience) at 4 °C for 30 min. APC fluorescence signals were determined using a Beckman flow cytometer and results were analyzed using GraphPad7 software.

### Pseudotype virus-based neutralization assay

SARS-CoV-2 pseudovirus encoding a luciferase reporter gene were purchased from NIFDC. HEK293T overexpressing hACE2 were seeded into the 96-well plates 16h before infection. 325 TCID50 pseudovirus were incubated with equal volume of 5 fold serially diluted antibodies overnight at 37 °C to reach equilibrium. The mixture of pseudoviruses and the tested molecules/Abs was added to the cells. After 24h incubation at 37 °C, cells were lysed and luminescence was measured with the Bio-Glo™ Luciferase Assay System (Promega). The IC50 values were calculated with non-linear regression log (inhibitor) vs. response (four parameters) using GraphPad Prism 7 (GraphPad Software, Inc.).

### Authentic virus plaque reduction neutralization test

Plaque reduction neutralization tests were done essentially as originally described^18^ with modifications^19^. Virus suspension (sequence-verified passage 4 of SARS-CoV-2 Victoria/01/2020) at appropriate concentrations in Dulbecco’s Modification of Eagle’s Medium containing 1 % FBS (D1; 100 μL) was mixed with test protein (antibody or recombinant chimeric protein, 100 μL) diluted in D1 at a final concentrations of 100 nM, 15.8 nM, 5 nM, 1.51 nM, 0.5 nM, 0.15 nM, 0.1 nM, and 0.018 nM, in triplicate, in wells of a 24-well tissue culture plate, and incubated at room temperature for 30 minutes. Thereafter, 0.5 mL of a single cell suspension of Vero E6 cells in D1 at 5 E5/mL was added, and incubated for 2 h at 37 °C before being overlain with 0.5 mL of D1 supplemented with carboxymethyl cellulose (1.5 %). Cultures were incubated for a further 4 days at 37 °C before plaques were revealed by staining the cell monolayers with amido black in acetic acid/methanol. The PRNT90 and PRNT50 values were calculated using the “FindECanything” non-linear curve fitting procedure of GraphPad Prism 8.4.1, with F set to 10 and 50, respectively.

## Acknowledgements

We appreciate the COVID-19 recovered patients’ blood samples from the Fifth Affiliated Hospital, Sun Yat-Sen University for the isolation and discovery of 89C8 antibody. The work in Oxford was supported by the generous support of philanthropic donors to the University of Oxford’s COVID-19 Research Response Fund”

## Supplementary material

**Supplementary Figure 1.**
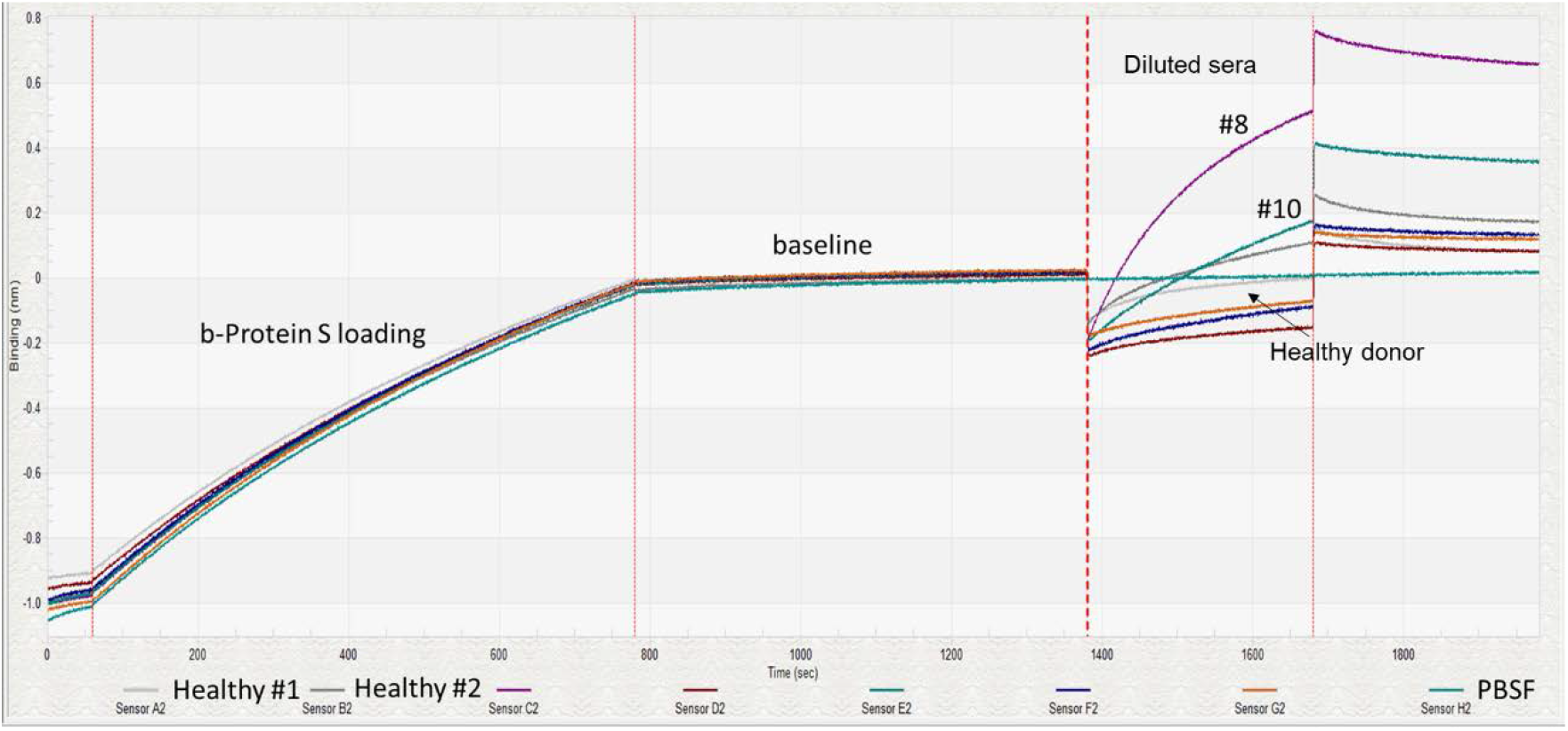
3 out of 10 sera samples from recovered COVID-19 patients showed stronger binding than healthy donor controls to biontinylated S1 on SA sensors by BLI.

**Supplementary Figure 2.**
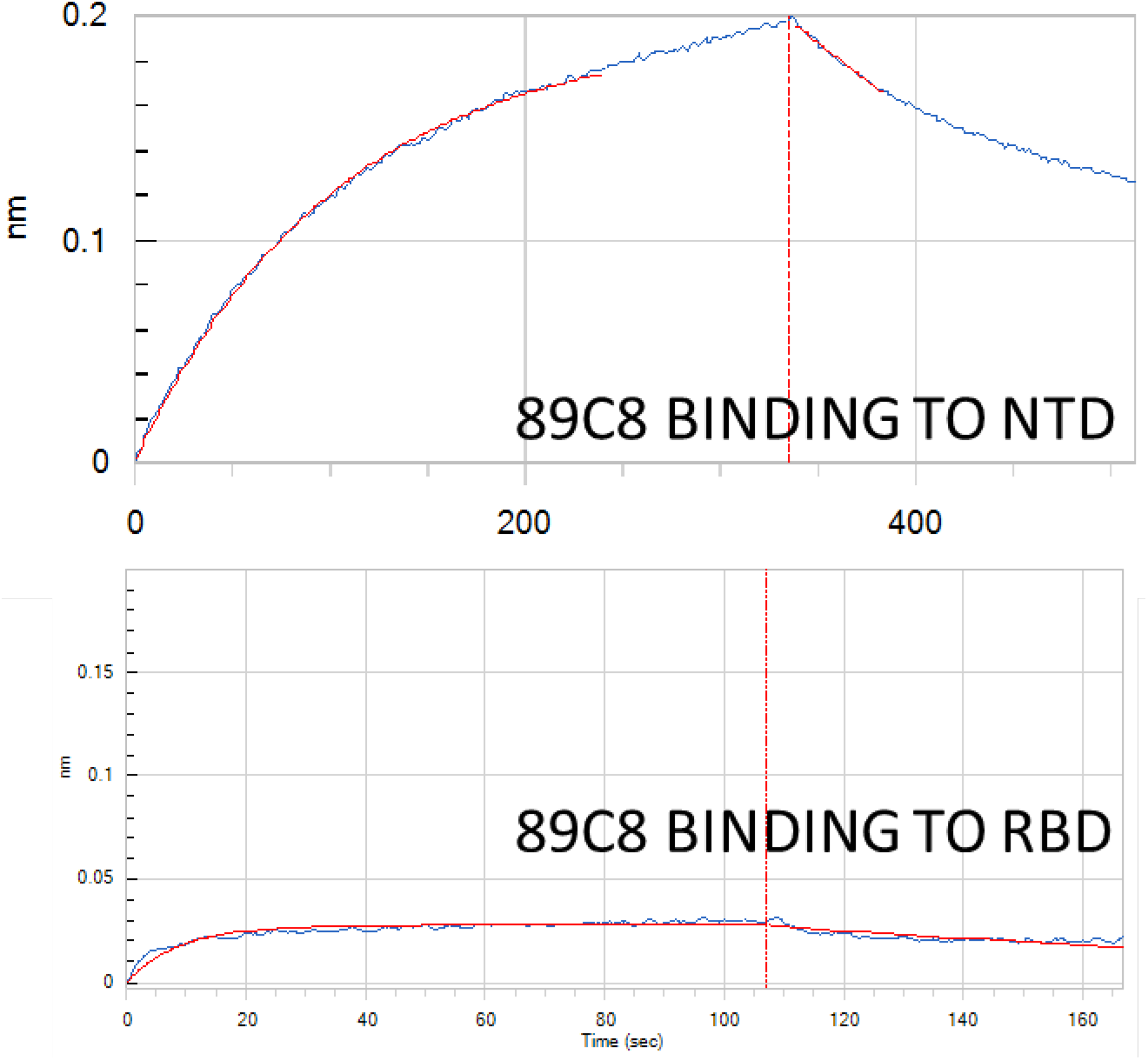
2a) 89C8 binding to NTD of S1 by BLI; 2b) 89C8 binding to RBD of S1 by BLI.

